# Geometric principles of second messenger dynamics in dendritic spines

**DOI:** 10.1101/444489

**Authors:** Andrea Cugno, Thomas M. Bartol, Terrence J. Sejnowski, Ravi Iyengar, Padmini Rangamani

## Abstract

Dendritic spines are small, bulbous protrusions along dendrites in neurons and play a critical role in synaptic transmission. Dendritic spines come in a variety of shapes that depend on their developmental state. Additionally, roughly 14*−*19% of mature spines have a specialized endoplasmic reticulum called the spine apparatus. How does the shape of a postsynaptic spine and its internal organization affect the spatio-temporal dynamics of short timescale signaling? Answers to this question are central to our understanding the initiation of synaptic transmission, learning, and memory formation. In this work, we investigated the effect of spine and spine apparatus size and shape on the spatio-temporal dynamics of second messengers using mathematical modeling using reaction-diffusion equations in idealized geometries (ellipsoids, spheres, and mushroom-shaped). Our analyses and simulations showed that in the short timescale, spine size and shape coupled with the spine apparatus geometries govern the spatiotemporal dynamics of second messengers. We show that the curvature of the geometries gives rise to *pseudo-harmonic* functions, which predict the locations of maximum and minimum concentrations along the spine head. Furthermore, we showed that the lifetime of the concentration gradient can be fine-tuned by localization of fluxes on the spine head and varying the relative curvatures and distances between the spine apparatus and the spine head. Thus, we have identified several key geometric determinants of how the spine head and spine apparatus may regulate the short timescale chemical dynamics of small molecules that control synaptic plasticity.

## 1 Introduction

Cell size, shape, and organelle location tightly regulate the dynamics of biochemical signal transduction; indeed even small molecule second messengers such as calcium (*Ca*^2+^), cyclic adenosine monophosphate (cAMP), and inositol trisphosphate (IP_3_) are reported to have distinct spatial microdomains within cells [1, 2]. Despite recent studies reporting the localization of these signaling molecules, the role of cell size and shape in controlling local intracellular signaling reactions, and how this spatial information originates and is propagated remains poorly understood. It has been hypothesized that spatial and temporal separation of second messengers can be a powerful means of specifying signaling functions through the interplay of cell shape and biochemical regulators [3, 4]. Therefore, an emerging concept in the understanding of signal transduction is that cell signaling is profoundly inhomogeneous in space, and that the spatio-temporal dynamics of signal molecules encode signaling specificity [5, 6]. This concept has been approached both theoretically [4, 7, 8] and experimentally [9–13].

One particular cell type where shape and signaling are closely related is the neuron. Communication in neurons is mediated by synapses and consists of complex signal transduction cascades. The presynaptic terminals release neurotransmitters that are then taken up by the post-synaptic spines to initiate a series of electrical, chemical, and mechanical events. Many of these events are tightly coupled to the dynamics of *Ca*^2+^, *cAMP*, and *IP*_3_ [14–16]. These second messengers are involved not only in the propagation of action potentials but also in downstream effects such as long-term potentiation (LTP), long-term depression (LTD), and structural plasticity. In particular, dendritic spines, which are thin post-synaptic protrusions, [17–19], have received much attention, especially because their density and morphology play a crucial role in mediating synaptic plasticity [20–24]. Changes in dendritic spine shape and density are symptomatic of several neuropathologies and neurodegenerative diseases such as Alzheimer’s, Parkinson’s, and drug addiction [25–29]. Thus, it is believed that the morphology of spines is closely related to their function: in fact, reciprocal changes between the structure and function of spines impact both local and global integration of signals within dendrites [22, 30–32]. Recently, Ramirez et al. [33] proposed that a combination of the geometry of dendritic spines and a characteristic propagation of Cdc42 induce localization of the protein in a timescale longer than the one estimated from the diffusion timescale alone.

Dendritic spines have characteristic shapes and internal organization; a schematic of the mushroom-like morphology of mature spines is shown in Figure 1. A very thin neck (40 ∼ 200 *nm* in diameter and 0.08 ∼ 1.1 *µm* long) separates an actin-rich bulbous head (0.3 ∼ 1.5 *µm* in diameter) from the dendrite, suggesting that the spine head acts as an isolated signaling compartment [6, 12, 13, 34–37]. Additionally, roughly 14*−*19% of spines contain a distinct organelle called the spine apparatus (SA), which is a protrusion of the smooth endoplasmic reticulum (ER). The spine apparatus may also be of various shapes that can change in response to signaling stimuli over days [6, 38–41]. It is generally accepted that changes in dendritic spine calcium levels, as well as localized protein synthesis, play a central role in structural plasticity, and these two processes may be influenced by the presence and shape of a spine apparatus. Experimental observations have shown that mice lacking spine apparatus (synaptopodin-deficient mice) show deficits in LTP and impaired spatial learning, thus supporting the hypothesis that the spine apparatus is involved in synaptic plasticity [42–48]. Furthermore, it has been hypothesized that computational modeling of the processes underlying structural plasticity could help to identify the regulatory feedback that governs the switch between LTD and LTP in ER-containing spines [49].

**Figure 1:**
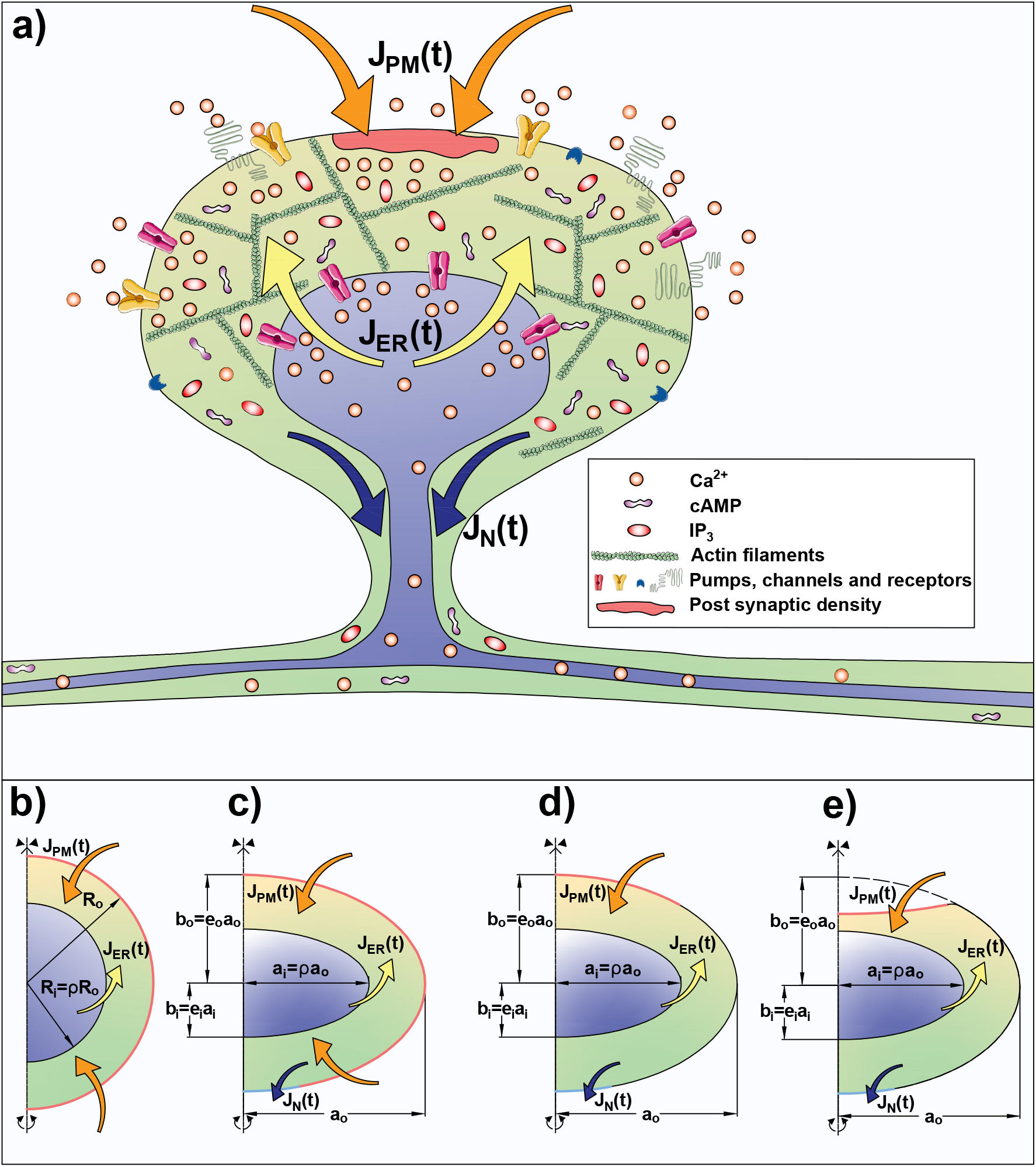
a) Schematic of a typical dendritic spine. Pumps, channels, and receptors on the plasma membrane (PM) and on the endoplasmic reticulum (ER) membrane allow for fluxes of second messengers such as *Ca*^2+^, *IP*_3_, and cAMP. These fluxes can be modeled as time-dependent flux boundary conditions, *J*_*PM*_ (*t*) at the plasma membrane and *J*_*ER*_(*t*) on the inner membrane. The effect of the presence of the neck has been included as an outlet flux *J*_*N*_ (*t*). Along with the reaction-diffusion dynamics in the domain, these fluxes determine the spatio-temporal dynamics of second messengers concentration in the dendritic spines. b-e) Geometries used to simulate the different spine shapes: b) spherical shell with outer radius *R*_*o*_ and inner radius *R*_*i*_ = *ρR*_*o*_ with uniformly distributed influx and no outlet; c) and d) oblate spheroidal shell with influx distributed throughout the head and localized on the pole of the spine respectively. The spheroids have outer eccentricity *e*_*o*_ and inner eccentricity *e*_*i*_. The dimensions of the outer shell are major axis *a*_*o*_ = *R*_*o*_ and minor axis *b*_*o*_ = *e*_*o*_*a*_*o*_ and the dimensions of the inner shell are major axis *a*_*i*_ = *ρa*_*o*_ and minor axis *b*_*i*_ = *e*_*i*_*a*_*i*_. e) Idealized mushroom-like geometry constructed by removing a portion of the shell in d).

Despite the emerging importance of the role of spine shape and internal organization in synaptic plasticity, the precise nature of how the physical aspects of a dendritic spine affect signaling dynamics of second messengers such as IP_3_, Ca^2+^, and cAMP remains poorly understood. In this paper, we conducted a systems biophysics study with the goal of identifying some of the design principles associated with the regulation of second messenger dynamics in dendritic spines. Specifically, using a combination of theory and computation, we sought to answer the following questions: (a) How are the spatio-temporal dynamics of second messengers within the spine affected by spine geometry? (b) How do the presence, size, and shape of the spine apparatus affect these spatio-temporal dynamics? And (c) how does different localization of the postsynaptic density (PSD, a protein-dense region in the postsynaptic membrane) affect the spatio-temporal dynamics of the second messenger? Using a minimal model to represent reaction-diffusion events within the spine, we showed that in the short timescale, the geometric features of the spine play an important role in establishing the spatio-temporal dynamics of the second messengers.

## 2 Methods

### 2.1 Governing equations

We developed and analyzed a reaction-diffusion model of second messenger dynamics with time-dependent flux boundary conditions. The resulting system of equations was analytically and numerically solved in simplified geometries to identify how the dynamics of second messengers are related to the geometrical parameters (see Figure 1b). We considered a second messenger with concentration distribution *C* = *C*(**x**, *t*), where **x** is the vector of the spatial coordinates and *t* is time. In the volume of the domain, the dynamics of *C* are then given by the following partial differential equation (PDE):

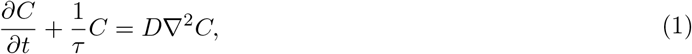

where *D* is the diffusion constant of the species *C*, *∇*^2^ represents the Laplacian operator in three dimensions, and *τ* is a time constant. *τ* represents a decay time constant associated with *C*. In the case of *Ca*^2+^, *τ* can be interpreted as the effective binding rate of rapid buffers. For *IP*_3_, *τ* represents the rate of degradation and for *cAMP* it represents the activity of phosphodiesterase. Our main goal in this study was to explore the solution to Eq. (1) for different geometries. Therefore, we chose a constant value of *τ* = 50 *ms*. Specialization of this model to *Ca*^2+^ and *cAM P* can be found in Bell *et al.* [50] and Ohadi *et al.* [51, 52], respectively.

### 2.2 Boundary conditions

To completely define the dynamics of *C* in the domain, we need to prescribe boundary conditions on both boundaries of the domain. Since the dynamics of the receptor-mediated events at the PM and on the organelle membranes are time-dependent fluxes due to signaling reactions [53–56], we prescribed the following boundary conditions

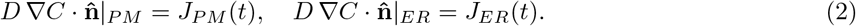

The fluxes *J*_*PM*_ and *J*_*ER*_ usually depend on many nonlinear reaction terms. For the purposes of our analyses, we used biexponential functions to describe the PM flux and included a slight delay in the ER flux to write

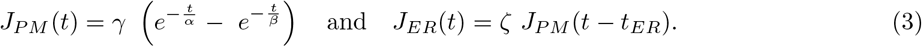

Here, the amplitude parameter *γ*, the time constants *α* and *β*, the amplitude of the PM-ER flux ratio *ζ*, and ER delay *t*_*ER*_ are free parameters that can be fit to experimental [57] or simulation data [6].

Finally, to simulate the effect of the efflux through the spine neck, we included an outlet flux in a portion of the outer membrane defined as

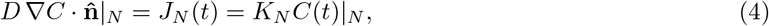

where *K*_*N*_ is a constant with units of a velocity (*µm/s*).

### 2.3 Geometries

We modeled the volume of the dendritic spine head using idealized geometries such that, to first approximation, they resemble the shape of a mature spine. We investigated three idealized geometries: spherical shells, oblate spheroidal shells, and idealized spheroidal mushroom-like geometries (see Figure 1b-e). The dimensions of the spherical shell shape are denoted as outer radius *R*_*o*_ and internal *R*_*i*_ = *ρR*_*o*_. To highlight the combined effect of curvatures and size of the membranes, we considered oblate spheroidal shells and spheroidal mushroom-like geometry with outer eccentricity *e*_*o*_ and inner eccentricity *e*_*i*_. As a result, the dimensions of the outer shell are major axis *a*_*o*_ and minor axis *b*_*o*_ = *e*_*o*_*a*_*o*_. The dimensions of the inner shell are major axis *a*_*i*_ = *ρa*_*o*_ and minor axis *b*_*i*_ = *e*_*i*_*a*_*i*_. The mushroom-like geometry was constructed by removing the intersection between the oblate spheroidal shell (Figure 1c) and another oblate spheroid centered at *b*_*o*_ in the vertical axis of symmetry with minor axis *b*_*o*_/10 and major axis *a*_*o*_ (Figure 1d). In this study, the geometrical parameters *ρ*, *e*_*i*_, and *e*_*o*_ were varied (in the range (0, 1]) to quantify the geometrical influence on the spatio-temporal dynamics of second messengers (*C*). For all simulations, we set the outer radius (and major axis) as *R*_*o*_ = *a*_*o*_ = 250 *nm*.

### 2.4 Computational tools

To assist in the derivation of the analytical solutions (Section 3.1), we used Mathematica 11.3 [58]. Simulations for the dynamics of second messengers were performed using finite element methods available through the commercial package COMSOL Multphysics 5.3a with MATLAB2018a [59]. In particular, the coefficient-form PDE interface was used along with parametrized geometries. The solutions were produced with the parametric sweep utility. Wolfram Mathematica 11.3 and MATLAB2018a [60] were used for post-processing the solutions and generating the figures.

### 2.5 Model analysis and relevant parameters

To compare the effect of different model geometries on the spatiotemporal dynamics of *C*, we defined the following metrics:

- **Gradient Metric**: In order to quantify a local measure of the spatial gradient of the species *C*, we defined

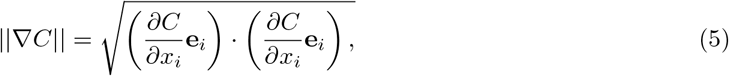

where *x*_*i*_ are the components of the position vector **x** in the direction defined by the vector bases **e**_*i*_, *i* = {1, 2, 3}, and the repeated indices indicate summation.
- **Extent of gradient**: A global measure of the variability of second messenger in the spine was defined as:

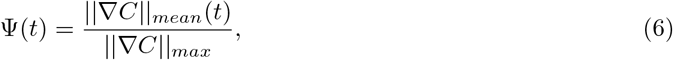

where ||∇*C*||_*mean*_(*t*) is the spatially averaged value of the gradient of *C* in Equation (5) and ||∇*C*||_*max*_ is the maximum peak value. This quantity gives us insight on the localization of second messengers. When Ψ(*t*) tends to zero, there is no concentration gradient in the spine.
- **Lifetime of the gradient**: We defined the lifetime of the gradient as the time, *τ*_Ψ_, needed to let Ψ(*t*) become lower than a threshold value Ψ_*th*_. In this work, we used Ψ_*th*_ = 25% as the value beyond which the distribution of *C* is uniform. This quantity gives us insight into the variation of the lifetime of the gradient with respect to different geometries and boundary conditions.

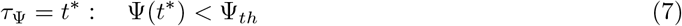

For clarity, the parameters used in the model are summarized in Table 1.

**Table 1:**
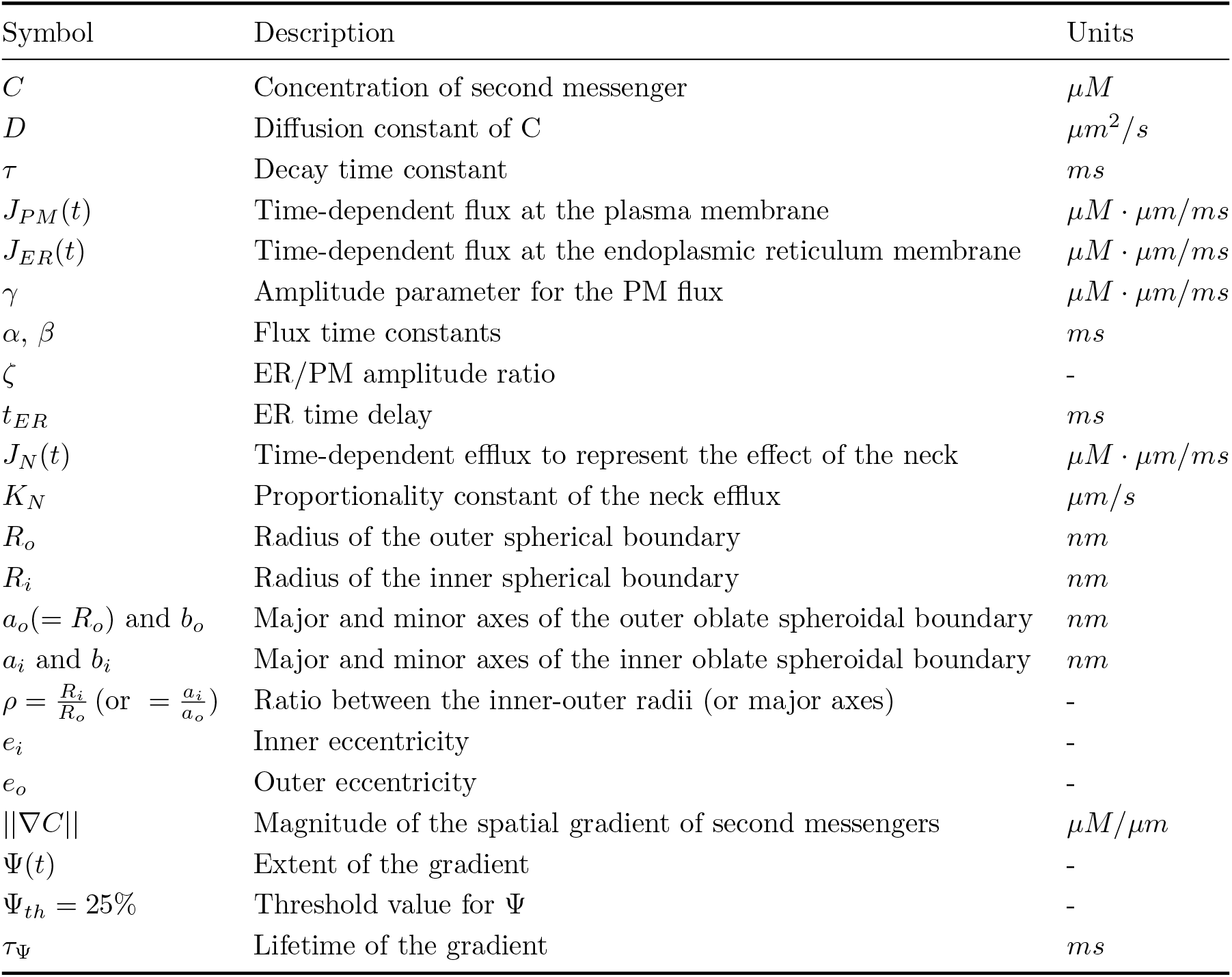
Notation used in the model

## 3 Results

### 3.1 Analytical solution

Equation (1) is a *homogeneous* PDE with time-dependent BCs given by Equation (2). To solve it, we used the method of *Generalized eigenfunction expansion* after formally transforming the problem into one with homogeneous boundary conditions; as a result, the PDE becomes nonhomogeneous [61]. We defined a function, the so-called *reference concentration distribution w*(**x**, *t*), such that it satisfies the given nonhomogeneous boundary condition, *i.e.*:

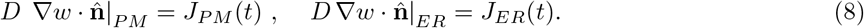

We considered a solution for *C* such that

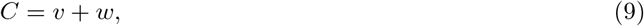

where the unknown function *v* is the solution of the new nonhomogeneous PDE with homogeneous BCs. This equation is obtained by substituting Equation (9) into Equation (1), resulting in,

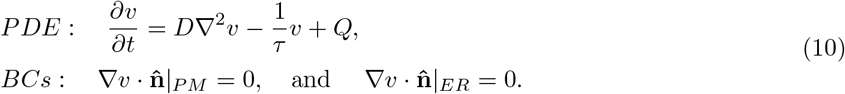

Here, 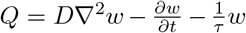 is a source term for *v*. For a homogeneous initial condition for *C*, the following initial condition holds for *v* and *w*,

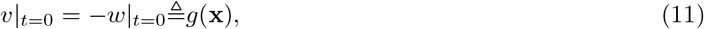

where *g*(**x**) is the initial condition for *v*. To solve Equation (10), the method of *eigenfunction expansion* is used, which consists of expanding the unknown function *v* in a series of *spatial* eigenfunctions Λ, resulting in

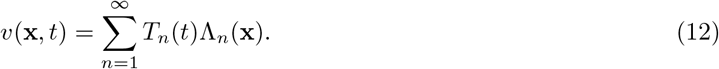

Here, *T*_*n*_(*t*) are the generalized Fourier coefficients of the eigenfunctions Λ_*n*_(**x**). The eigenfunctions can be found using the *associated* homogeneous version (*Q* = 0) of Equation (10). Using the method of separation of variables, *v* = *T* (*t*)Λ(**x**), we can write,

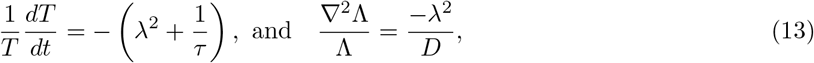

where *λ* is a separation constant.

The separation between time and space highlights the *pseudo-harmonic* nature of the distribution of second messenger (*C*). The temporal ordinary differential equation (ODE in Equation (13)) has an exponential decay as the solution, which is affected by both *λ* and the timescale *τ*. The spatial PDE in Equation (13) represents the Helmholtz wave equation, which is separable in 11 three-dimensional coordinate systems, and for each of them, there exists a specific family of solutions, known as the harmonic wave functions [62, 63]. However, the presence of internal and external time-dependent boundary conditions and the transcendental nature of the kernel of the solution limit the possibility of finding explicit solutions and restrict the solution to an implicit form. In what follows, we further specialize this analysis for spherical domains with and without an inner boundary (Sections 3.1.1 and 3.1.2) and oblate spheroidal shells (Section 3.1.3).

#### 3.1.1 Analytical solution in spherical shells with uniformly distributed BCs

We first considered a spherical shell with an inner boundary of radius *R*_*i*_, an outer radius of *R*_*o*_, and with uniformly distributed influx on both membranes (see Figure 1c). We used a spherical coordinate system (**x** = (*r* **e**_**r**_, *θ* **e**_*θ*_, *φ* **e**_*φ*_)) defined in terms of the Cartesian (*x, y, z*) coordinates as shown in Figure 2a, where *r ∈* [0, *∞*), *θ ∈* [0, 2*π*), and *φ ∈* [0, *π*] represent the radial, the azimuthal, and polar coordinates respectively [65]. Exploiting the spherical symmetry of both the domain and the boundary condition, we assumed that *C*(**x**, *t*), and thus *v*(**x**, *t*) and *w*(**x**, *t*), depend only on the radial coordinate *r*. That is, we assume that there is no dependence on the angular coordinates *θ* and *φ*. Thus, the Laplacian and normal gradient operators are:

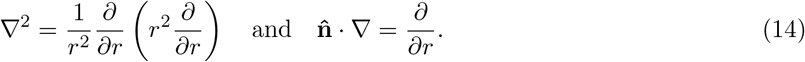

**Figure 2:**
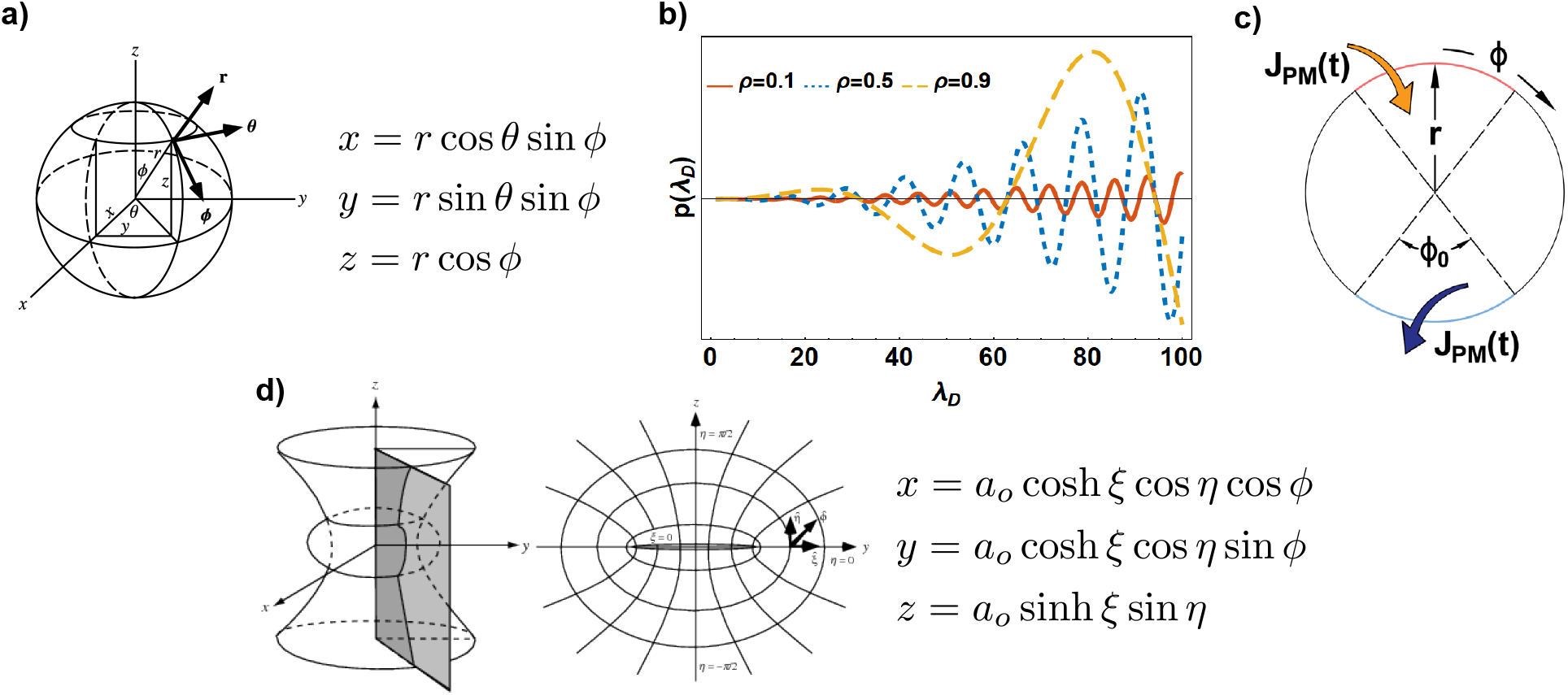
a) 3D representation and definition of the spherical coordinate system [64]; b) Function *p*(*λ*_*d*_) Equation (20) for different values of the ratio *ρ*. All of the zero-crossings represent the eigenvalues *λ*_*n*_ of Equation (13); c) Spherical domain with anti-periodic BCs to study both influx and efflux BCs; d) 3D representation, 2D representation in the y-z plane, and definition of the oblate spheroidal coordinates system [65].

In this framework, the simplest way to define *w*(*r, t*) respecting the conditions in Equation (8) is

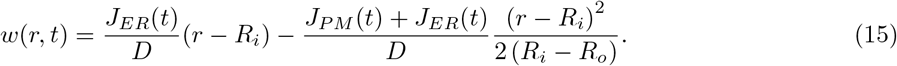

The homogeneous initial condition for *C* and Equation (15), result in the initial condition for *v* as,

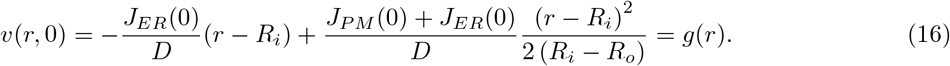

The solution of the Helmholtz Equation (Equation (13)) in spherical coordinates is the sum of spherical Bessel functions of the first kind (*j*) and the second kind (*y*) of zero order [61–63],

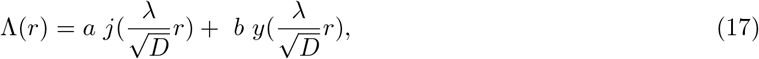

where *a* and *b* are integration constants. Due to the homogeneity of both the PDE and BCs, there exists a non-trivial solution for *v* if we impose the *determinant condition* [61]. The BCs in Equation (10) now become

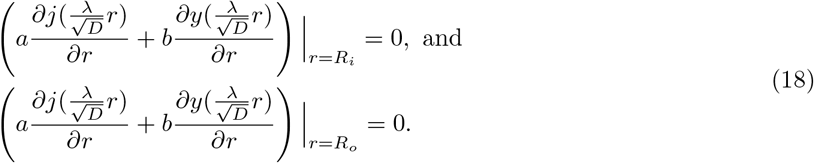

The eigenvalues *λ*_*n*_ are the roots of the *characteristic polynomial*

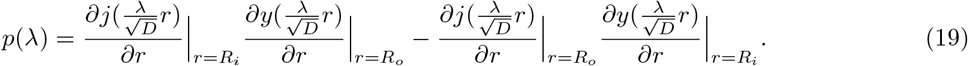

After some algebraic manipulation and exploiting the properties of the Bessel functions, it is possible to rearrange *p*(*λ*) as

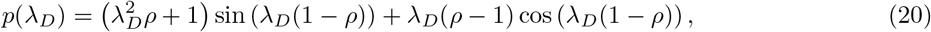

where 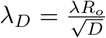 and 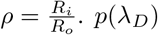 is a transcendental function and the zeros can be found only numerically as showing in Figure 2b.

Assuming that the eigenvalues *λ*_*n*_ have been found, the expansion in Equation (12) can be specialized as follows,

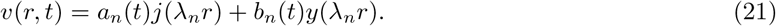

Exploiting the orthogonality of the eigenfunctions, we can now determine the initial values of the generalized Fourier coefficients as [61]

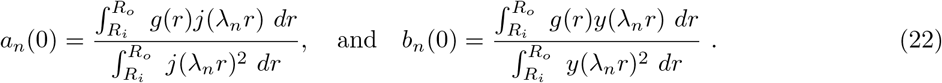

Substituting Equation (21) back into Equation (10), we obtain two nonhomogeneous first order ODEs for the Fourier coefficients that must respect the initial conditions in Equation (22),

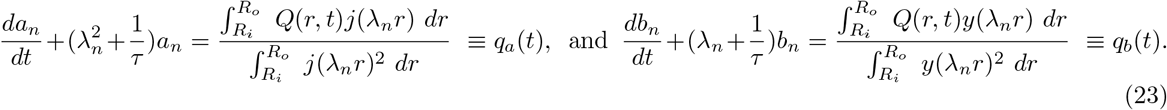

The solutions of the above equations are given by

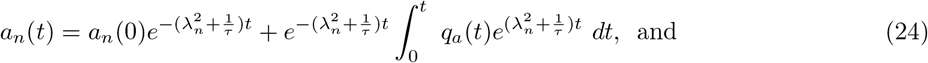

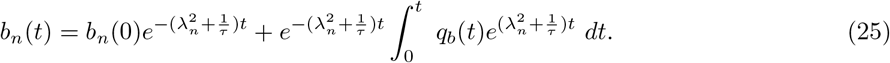

Despite the complexity of these coefficients, we observed that the amplitude of the wave functions in Equations (24) and (25) are implicitly related to the temporal dynamics of the biochemical reactions on the membranes of the spine (through *Q*), intricately coupled by the morphology (through *λ*_*n*_), and reaction kinetics (through *τ*).

#### 3.1.2 Analytical solution in spherical domains with anti-periodic BCs

We next analyzed the effect of localized influx and efflux at the outer membrane representative of fluxes at PSD and at the neck, respectively. In this context, we investigated the analytical solution of Equation (1) in a spherical domain with no inner boundary and anti-periodic BCs at the outer domain (see Figure 2c) written as

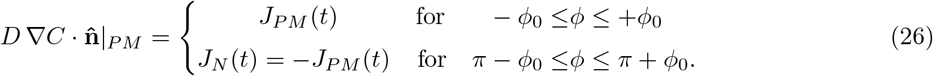

Here, *J*_*N*_ reflects efflux through the neck. We assumed that the amplitude of the influx and efflux are the same to simplify our analysis and obtain a semi-analytical solution by exploiting the axial symmetry of the domain and BCs. As a result, *C*(**x**, *t*) and therefore *v*(**x**, *t*), and *w*(**x**, *t*) depend on *r* and *φ* but not *θ*. The Laplacian and normal gradient operators in spherical coordinates (Figure 2a) are

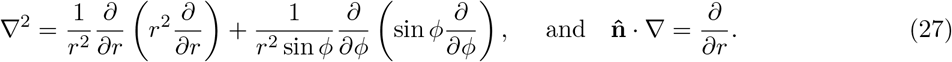

In this framework, the simplest way to define *w*(*r, φ, t*) respecting Equation (8) and Equation (26) is

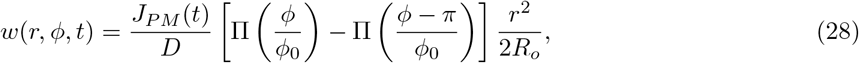

where Π(*φ*) is a rectangle function. Given the homogeneous initial condition for *C* and Equation (28), the initial condition for *v* is

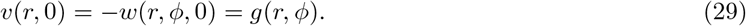

Using separation of variables, Λ = *R*(*r*)Φ(*φ*), the Helmholtz equation in Equation (13) leads to the following two equations

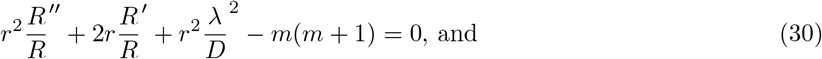

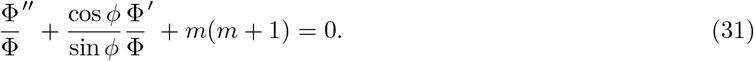

Here, *m* is a separation constant. Due to the anti-symmetric periodic condition (Φ[0] = *−*Φ[*π*]), the solution of Equation (31) involves the Legendre functions, *L*_*m*_(*φ*), of the first kind and of odd *m*^*th*^ order,

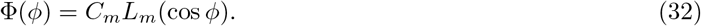

The solutions of Equation (31) are spherical Bessel functions of the first kind of *m*^*th*^ order (*j*_*m*_(*r*)) [61–63, 65],

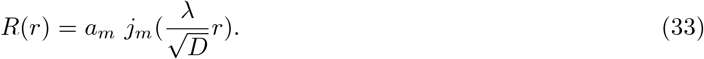

To find the *λ*_*mn*_ eigenvalues we need to enforce homogeneous BCs 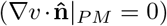 that correspond to finding the roots of derivative of the Bessel functions such that

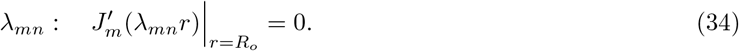

Assuming that the eigenvalues *λ*_*mn*_ can be found, the expansion in Equation (12) can be written as

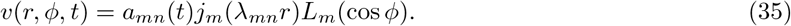

Exploiting the orthogonality of the eigenfunctions the initial values of the generalized Fourier coefficients are given by

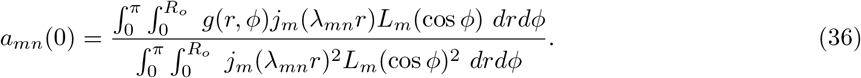

Substituting Equation (35) back into Equation (10), we obtain a nonhomogeneous first order ODE for the Fourier coefficients that must respect the initial conditions in Equation (36),

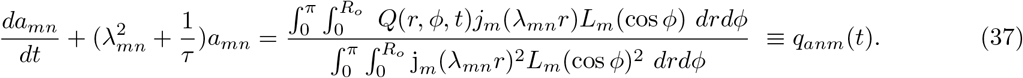

The solution of the above equation leads to the following expression for *a*_*mn*_(*t*),

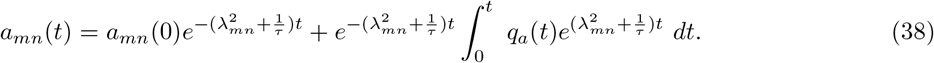

It is worth noticing that, with respect to the uniformly distributed BCs, the asymmetry of the localized fluxes introduced new angular harmonics *L*_*m*_ in addition to the radial wave functions *J*_*m*_. Thus, in addition to the complex coupling between morphology of the spine and degradation kinetics, we found that the spatial distribution of the biochemical reactions on the membrane of the spines (through *φ*_0_) also regulates the dynamics of second messengers.

#### 3.1.3 Analytical solution in oblate spheroidal shells with uniformly distributed BCs

We next obtained the analytical solutions for the case of the confocal oblate spheroidal shell with uniformly distributed BCs (Equation (2)). The dimensions of the outer shell are major axis *a*_*o*_ and minor axis *b*_*o*_ = *e*_*o*_*a*_*o*_.

The dimensions of the inner shell are major axis *a*_*i*_ = *ρa*_*o*_ and minor axis *b*_*i*_ = *e*_*i*_*a*_*i*_. We used the oblate spheroidal coordinates system **x** = (*ξ* **e**_*ξ*_, η **e**_*η*_, φ **e**_*φ*_) related to the Cartesian coordinates as shown in Figure 2d where *ξ ∈* [0, *∞*), *η ∈* [*−π*/2, *π*/2], and *φ ∈* [0, 2*π*). Surfaces with constant *ξ*, *η*, and *φ* are oblate spheroids, confocal one-sheeted hyperboloids of revolution, and half-planes through the z-axis, respectively (Figure 2d). To express the Laplacian in oblate spheroidal coordinates, an alternative set of spheroidal coordinates, *ε*_1_ = sinh *ξ*, *ε*_2_ = sin *η*, and *ε*_3_ = *φ*, can be used ([62, 63, 65–69]) as follows

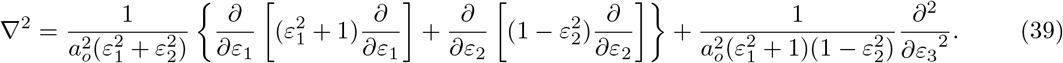

The normal gradient to the boundary of oblate ellipsoids is given by

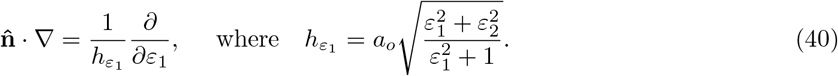

We assume that it is possible to establish a reference function *w* such that the BCs (Equation (8)) in oblate spheroidal coordinates Equation (40) are satisfied. In this framework, the Helmholtz equation in Equation (13) has solution in the form Λ = *w*_1_(*ε*_1_)*w*_2_(*ε*_2_)*w*_3_(*ε*_3_), where *w*_1_, *w*_2_, and *w*_3_ satisfy the following *spheroidal wave* equations respectively [62, 63, 66–69],

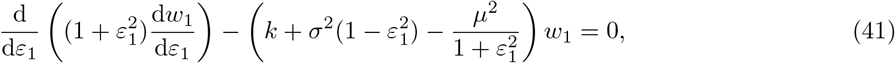

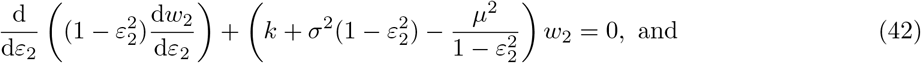

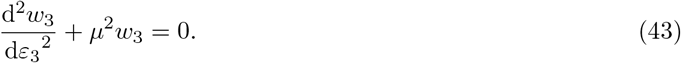

with 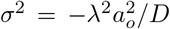 and, *k* and *µ*^2^ are new separation constants. The solutions of Equations (41)–(43) involve the *spheroidal harmonic functions*

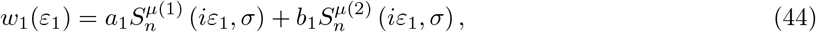

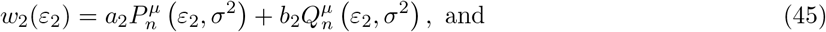

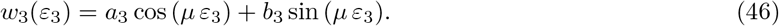

Here, the eigenfunctions *S*, *P* and *Q* are the so-called *radial wave* functions, *spheroidal wave functions of first kind*, and *spheroidal wave functions of second kind* respectively, while *a*_1_, *a*_2_, *b*_1_, *b*_2_, *c*_1_, and *c*_2_ are integration constants. The oblate spheroidal solution can be simplified assuming the oblate symmetry of both the geometry and the BCs and thus we write Λ = *w*_1_(*ε*_1_)*w*_2_(*ε*_2_) and *µ* = 0. Also in this case, the associated eigenvalues can be found by imposing the determinant condition in an implicit form. Similar to the spherical case (Equations (24)–(25)) the generalized Fourier coefficients (Equation (12)) will have an exponential time decay highlighting the pseudo-harmonic nature of the solution. As a result a deviation from the spherical shape introduces more complex spatial dependence for *C*. In addition to the radial variation, new angular wave functions regulate the spatiotemporal dynamics of second messengers in the *φ* direction.

### 3.2 Combined effect of spine apparatus size and diffusion coefficient on the lifetime of second messengers gradient

Thus far we have elaborated on the pseudo-harmonic nature of the solution showing how the analytical solutions can be provided implicitly. In order to visualize these solutions, we now explore the parameter space numerically. We begin with a simulation of the spatiotemporal dynamics of *C* in a spherical spine head with a radius *r* = *R*_*o*_ and a spherical spine apparatus with a radius *R*_*i*_ = *ρR*_*o*_ (Figure 1 and Table 1). For simplicity, we start with uniformly distributed influx (*J*_*PM*_ (*t*) and *J*_*ER*_(*t*)) boundary conditions on the outer and inner boundary but with no outlet (*J*_*N*_ (*t*) = 0). For illustrative purposes, we chose a set of parameters for the boundary conditions (Figure 3a) such that we can match previously reported dynamics of calcium [6]. For a diffusion coefficient *D* = 100 *µm*^2^/*s* and *ρ* = 0.5, the profile of *C* at a location exactly halfway between the two membranes is shown in (red line, Figure 3b); this temporal profile is in good agreement with previous observations (see inset in Figure 3b). Therefore, for all the following simulations, we used the time constants (*α, β*, and *t*_*ER*_) as shown in Figure 3a. The amplitude parameters (*γ* and *ζ*) have been customized to the specific geometry, to avoid abnormal peaks of concentration of second messengers.

**Figure 3:**
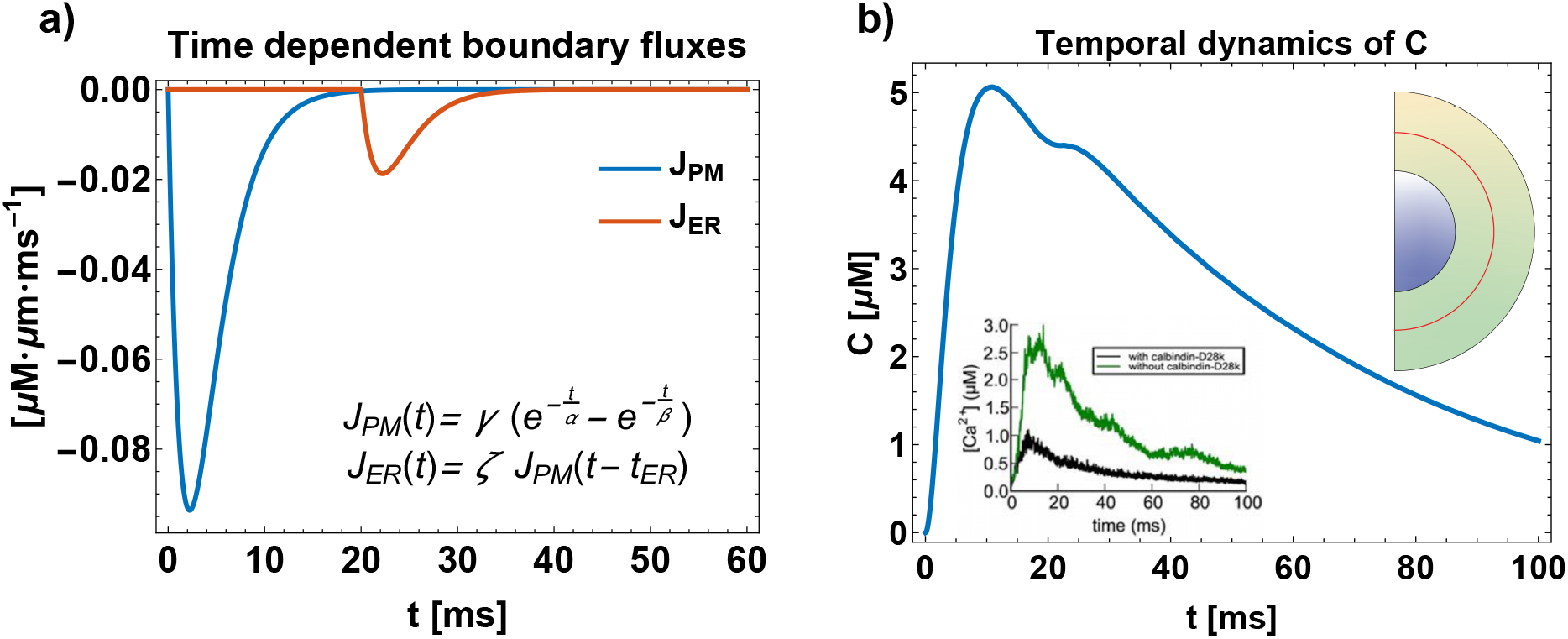
a) Time-dependent boundary fluxes for the outer membrane (solid blue) and inner membrane (dashed red) respectively, representing the dynamics of various pumps and channels; *D* = 100 *µm*^2^/*s*, and *ρ* = 0.5. The flux parameters (*α* = 2.5 *ms, β* = 2 *ms, γ* = *−*1.14 *×* 10^*−*6^ *µM ⋅ µm/ms, ζ* = 0.2, and *t*_*ER*_ = 20 *ms*) have been fitted to reproduce typical temporal dynamics of second messengers inside a dendritic spine. b) Temporal dynamics of *C*, in *µM*, at the midpoint between the inner and outer shells (*r\** = (*R*_*o*_ − *R*_*i*_)/2 + *R*_*i*_, red line). Inset shows the data from MCell simulation for *Ca*^2+^ dynamics [6].

A characteristic feature of the spatio-temporal dynamics of *C* is the lifetime of the gradient, which is affected by both the ratio between the radii (*ρ*) and the diffusion coefficient (*D*). To decode how these two quantities affect the spatiotemporal dynamics of *C*, we conducted the following simulations – (i) spine apparatus size was varied by changing *ρ*; we used three different values of *ρ* (0.1, 0.5, and 0.9) to capture the extreme volume changes due to small, medium, and large spine apparatus. (ii) The diffusion constant of *C* was varied to capture the range of intracellular diffusion from a crowded regime to free diffusion (1, 10, 100 *µm*^2^/*s*) [70–76]. We found that with small spine apparatus (*ρ* = 0.1), a significant concentration gradient exists in the radial direction when the diffusion coefficient is small (*D* = 1*µm*^2^/*s*) at early time points but not for larger diffusion coefficients (Figure 4a). As the spine apparatus gets larger, there is less volume available in the spine and more surface area for the internal membrane and consequently more flux through the ER. Therefore, the variability of *C* is smaller even for a lower diffusion coefficient (Figure 4a-c).

**Figure 4:**
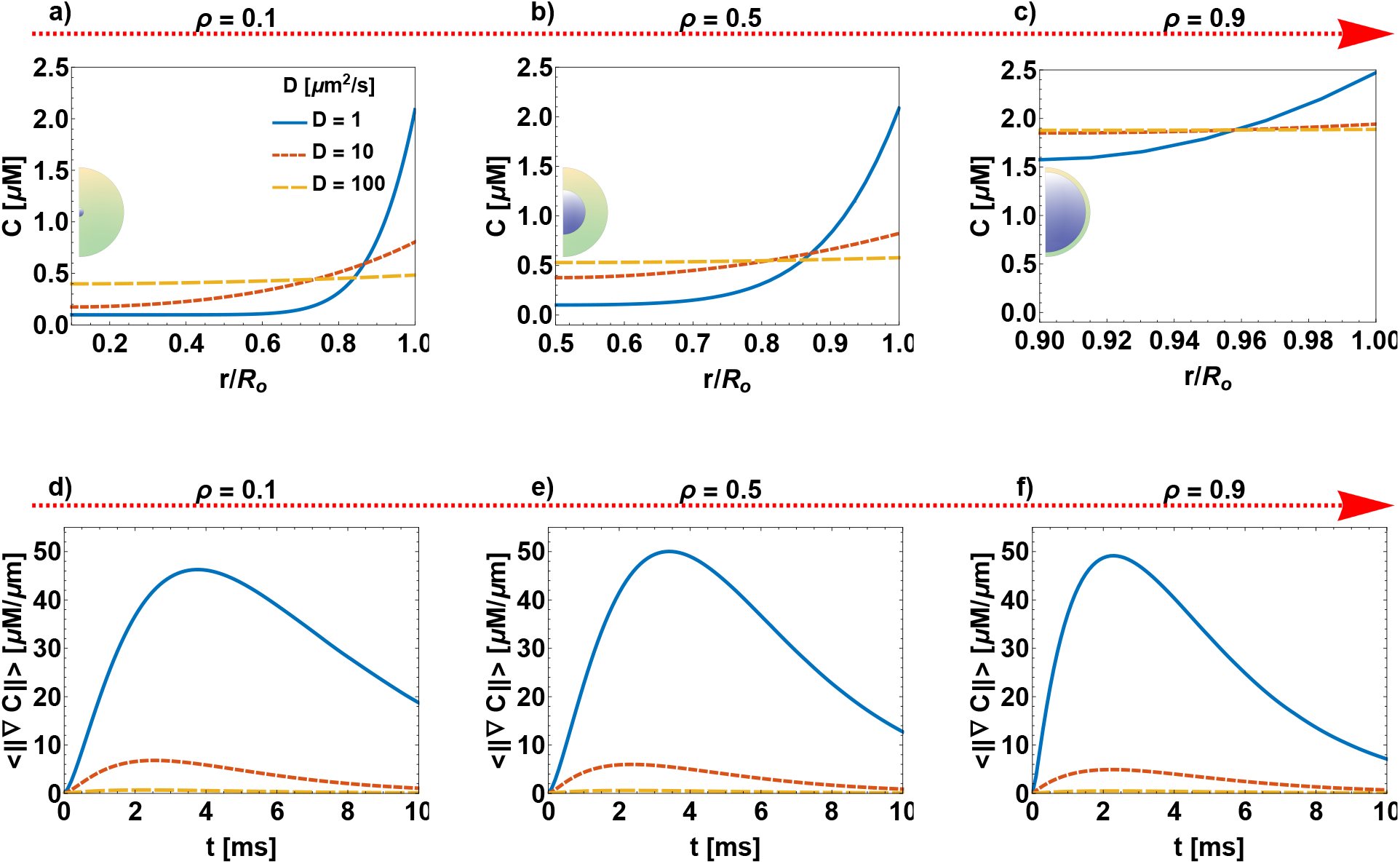
Effect of spine apparatus size and diffusion constant *D* on the lifetime of *C*. The top row shows the radial distribution of *C* concentration for three different values of diffusivities (*D* = 1, 10, 100 *μm*^2^/*s*) at 1 *ms*: a) *ρ* = 0.1, b) *ρ* = 0.5, and c) *ρ* = 0.9. The bottom row shows the time course of the mean value in the domain of the concentration gradient (*< ||∇C|| >*), for d) *ρ* = 0.1, e) *ρ* = 0.5, and f) *ρ* = 0.9.

To quantify the lifetime of the gradient, we calculated the norm of the gradient of *C* (Equation (5)) as a function of time. Again, when the spine apparatus is small and the diffusion coefficient is small, the gradient lasts longer. An increase in diffusion coefficient for a given spine apparatus size reduces the lifetime (Figure 4d). As the size of the spine apparatus increases, even for small diffusion coefficients, the lifetime of the gradient decreases (Figure 4d-f) confirming that small spine apparatus and low diffusion will result in longer time gradients of *C*. On the other hand, a large apparatus, even with a small diffusion coefficient will result in rapid propagation of second messengers from the PM to the ER membrane.

Given that our results are sensitive to the value of diffusion coefficients, we surveyed the literature for estimates of diffusion coefficient of different second messengers (Table 2). Based on this survey and the fact that the spine has a highly crowed environment characterized by a viscosity 5 times higher than the other cell types [77], we chose a value of *D* = 10 *μm*^2^/*s* for the remaining simulations.

**Table 2:** 20-30 fold slow down for the diffusion constant of second messengers in dendritic spines

| Species | D [$\mu m^2/s$] | Method | Ref. |
| --- | --- | --- | --- |
| $cAMP$ | [5 – 76] | experimental | Agarwal *et al.* [70] |
|  | 60 | simulations | Yang *et al.* [71] |
| $IP_3$ | [3 – 10] | experimental | Dickinson *et al.* [72] |
| $Ca^{2+}$ | [13 – 65] | experimental | Allbritton *et al.* [73] |
|  | < 16 | experimental | Gabso and Neher [74] |
|  | 30 | experimental & simulation | Biess *et al.* [75] |
|  | 1.5 | theory | Matthews and Dietrich [76] |
|  | 7 | theory (Stokes–Einstein equation) | Cai *et al.* [78] |

### 3.3 Size and shape of the spine and spine apparatus affect the gradient of signaling molecules

We next investigated the effect of the shape of the spine and the spine apparatus. This is particularly relevant since dendritic spines are known to have distinct shapes and their shape is associated with function [22, 30, 31]. Because the shape of the spine is a result of many different geometric properties, we focused specifically on curvature by modeling the spine shapes as different ellipsoidal shells. Even though this is a mathematical idealization, the resulting solutions provide insight into how curvature variations along the ellipsoids affect the harmonic functions that govern the profile of *C*, for uniformly spatially distributed boundary conditions. As before, the ratio between PM-ER size is controlled by *ρ* and now the shape is controlled by the eccentricities of the inner and outer ellipsoids *e*_*i*_ and *e*_*o*_ respectively (Figure 1c and Table 1). We conducted a systematic variation of the magnitude of these geometrical parameters and analyzed their effect on the spatio-temporal dynamics of *C* (Figure 5). The influx *J*_*PM*_ (*t*) was considered distributed on all the outer boundary except for a smaller portion (*⋍* 0.2 *μm* in diameter [12]) to include the presence of an outlet flux due to the neck *J*_*N*_ (*t*) (see Figure 1c).

**Figure 5:**
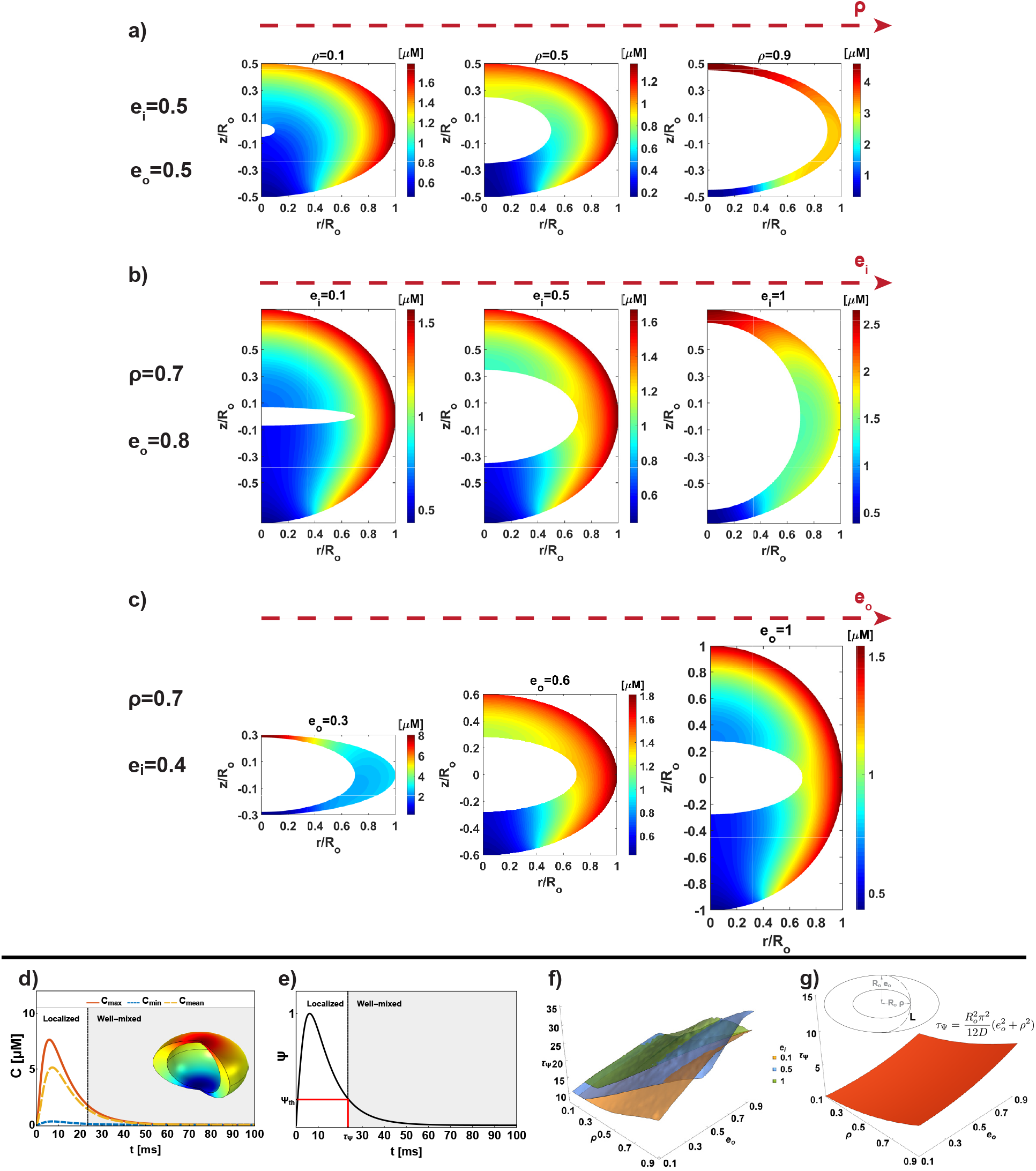
Effect of spine shape and spine apparatus shape on the spatial distribution of *C*. The influx *J*_*PM*_ (*t*) is distributed on all the outer boundary except for a small portion (*⋍* 0.2 *μm* in diameter) to include the presence of an outlet flux *J*_*N*_ (*t*) on the neck. All simulations are shown for *D* = 10 *μm*^2^/*s*, *R*_*o*_ = 250 *nm,t* = 1 *ms*, and *K*_*N*_ = 1 *μm/s*. a) For *e*_*i*_ = 0.5 and *e*_*o*_ = 0.5, we analyzed the effect of increasing *ρ*. As *ρ* increases, the location of maximum concentration changes from the equator to the pole, and the maximum value increases as well. Similar behavior is highlighted in b) where the effect of the increase in internal eccentricity is shown, kipping *ρ* = 0.7 and *e*_*o*_ = 0.8 constant. In comparison, c) shows opposite behavior, when moving toward a more spheroidal outer shape (*e_o_ →* 1), maintaining fixed *ρ* = 0.7 and *e*_*i*_ = 0.4. For the geometrical parameters *ρ* = 0.7, *e*_*i*_ = 0.8, and *e*_*o*_ = 0.6, we plotted the time evolution of *C*_*max*_, *C*_*min*_, and *C*_*mean*_ in d). A significant localization holds for few ms before a well-mixed distribution is reached. The extent of gradient Ψ is plotted in e) and we showed in f) the surfaces *τ*_Ψ_ as a function of *ρ*, *e*_*i*_, and *e*_*o*_ highlighting how the lifetime gradient nonlinearly depends on the geometrical parameters. The theoretical lifetime of the gradient is shown in g) considering a characteristic length *L* illustrated in the inset. The lifetimes from theory and numerical simulations follow similar nonlinear dependence to the geometrical parameters (*ρ*, and *e*_*o*_). Note that the lifetime from simulation includes the coupling among diffusion, geometry, timescale of the degradation (*τ*), timescale of boundary conditions (*α* and *β*), as well as their localization whereas the analysis is based on length scale and diffusivity alone.

For a given value of *e*_*o*_ and *e*_*i*_, both set to 0.5 in this case, as *ρ* increases, the location of maximum concentration changes from the equator to the pole. This is because of the *pseudoharmonic* nature of the solutions and a complex interplay among curvatures and distance between the membranes that induces a nonuniform distribution of the surface (and thus flux) per available volume (Figure 5a). Where the outer membrane is more curved, there is more surface per volume and this is where higher levels of concentration are attained. The increase in flux per volume can also be caused by changing the distance between the membranes. In fact, when this distance is too small, the curvature effects became secondary. Furthermore, when there is less volume available in the spine (increasing *ρ*), the overall concentration of *C* increases. For a constant *ρ* and *e*_*o*_, increasing *e*_*i*_, which results in the spine apparatus tending towards a spherical geometry (Figure 5b), *C* passes from a spherical-like distribution (*e*_*i*_ = 0.1) to a localization of the maximum on the pole (*e*_*i*_ = 1) through an intermediate situation where the maximum is confined at the equator (*e*_*i*_ = 0.5). These results hold even in the case of constant *ρ* and *e*_*i*_ and decreasing *e*_*o*_ (Figure 5c). Thus, the shape and size effects of the spine and spine apparatus are a result of PM shape and ER membrane shape and the relative volume enclosed. In fact, a significant difference between maximum and minimum peaks hold for tens of milliseconds, before a well-mixed condition is reached (Figure 5d), as showed by the asymptotic evolution of the extent of gradient Ψ (Figure 5e).

We plotted *τ*_Ψ_ (Table 1) for three constant values of *e*_*i*_ (0.1, 0.5, and 1) as a function of *ρ* ([0.02, 0.9]) and *e*_*o*_ ([0.1, 0.9]) (Figure 5f). If *τ*_Ψ_ were unaffected by geometrical parameters, we would expect to see a flat plane for each value of *e*_*i*_. We observed that the lifetime of the gradient depends on *ρ* and *e*_*o*_ in a nonlinear manner and it is not affected by the inner eccentricity. A big ER in a spheroidal*−*like spine (*ρ →* 1 and *e_o_ →* 1) represents a geometrical barrier against the diffusion of *C* (*τ*_Ψ_ *⋍* 35 *ms*). On the contrary, a spine with a flattened head (small *e*_*o*_) with a very small or absent ER (*ρ →* 0) has a 4-fold shorter lifetime of the gradient (*τ*_Ψ_ *⋍* 8 *ms*). We found that the paraboloidal trend of *τ*_Ψ_ from simulation is in very good agreement with the one from theory, *τ*_Ψ_ *⋍ L*^2^/6*D*. Here, the length *L* is defined as the semi-perimeter of the ellipses with axis *a*_*i*_ = *ρR*_*o*_ and *b*_*o*_ = *e*_*o*_*R*_*o*_ and calculated with the approximated formula 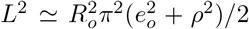 (see the inset in Figure 5g). It is worth highlighting that the simulations include coupling with the timescale of the reaction, the dynamics due to the boundary conditions, as well as their distribution at the boundary of the domain whereas the analysis is based on length scale and diffusivity alone. This results in an overall extension of the lifetime of the gradient with respect to the theoretical estimation.

The results from these simulations can be summarized as follows: *first*, unlike spherical shells where the spatial variation is only in the radial direction, ellipsoidal shells show a spatial variation in the *z* and the *r* directions (Figure 5a-c); *second*, the spatial variation of *C*, particularly the location of high and low concentrations of *C* at a given time can switch from the equator to the pole or *vice versa* depending on the geometry alone (Figure 5a-c); *third*, the nonlinear trend of the timescale *τ*_Ψ_ can be understood in terms of classical diffusion lifetime as long as the length scales are corrected for the geometries (Figure 5d-g). Thus, these simulations predict that a deviation from spherical shape provides spines access to a more complex phase space with respect to the properties of the gradient of *C* and its lifetime.

### 3.4 Consequences of a localized input of second messengers in dendritic spines

Thus far, we have shown that geometry by itself, in the presence of uniformly distributed boundary conditions, produces transient localization of second messengers. However, in reality, the influx at the outer boundary is not uniformly distributed but is mostly localized to specific regions of the spine head. In fact, as shown by the reconstruction of many dendritic spines [6, 12, 13, 40, 79–82], the post-synaptic density, a portion of the PM rich in ionotropic receptors is localized along different positions of the membrane. Furthermore, the spine neck acts as a sink for second messengers, allowing signal transmission toward the dendritic shaft. Therefore, we next analyzed the combined effect of spine geometries, influx BCs localized to specific portions of the boundary and presence of an outlet in the neck region (Figure 6).

**Figure 6:**
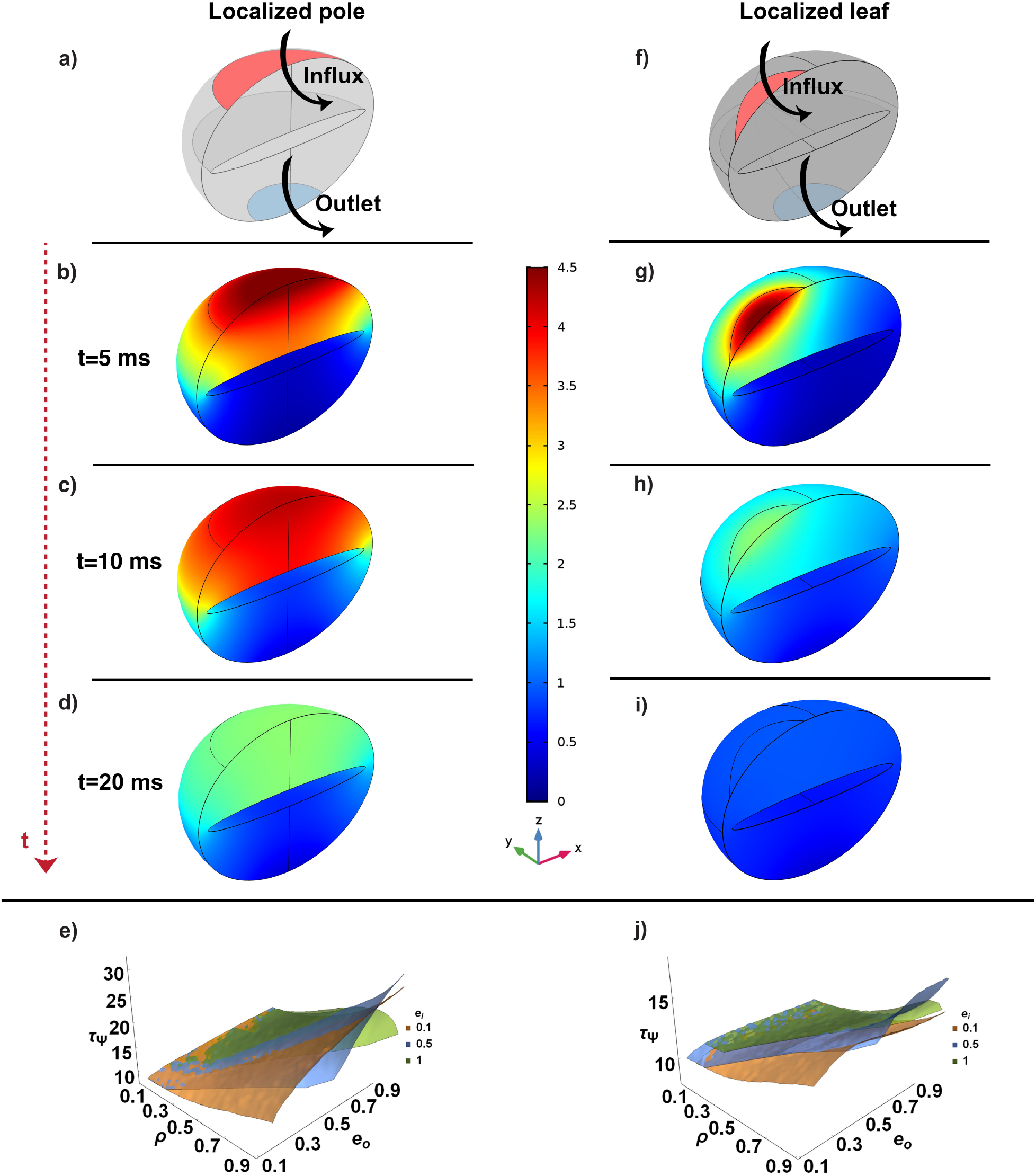
Effect of localized flux at the outer membrane of the spine. All the simulations shown use *D* = 10 *μm*^2^/*s*, *R*_*o*_ = 250 *nm*, and *K*_*N*_ = 1 *μm/s*. In the schematic a) and f) we analyzed the localized pole and leaf influx respectively, both with an outlet, considering the geometrical parameters *ρ* = 0.9, *e*_*o*_ = 0.8, and *e*_*i*_ = 0.1. We showed the distribution of *C* at *t* = 5 ms b) and f), *t* = 10 ms c) and f), and *t* = 20 ms d) and e). *τ*_Ψ_ from the simulations for *e*_*i*_ = 0.1, 0.5, and 1 for the pole and leaf case are shown in e) and j) respectively. The geometrical barrier represented by a big ER (*ρ →* 0.9) leads to a much more persistent localization (*τ*_Ψ_ = [10 ∼ 30] *ms*) but the localization of the influx at the side reduces this effect *τ*_Ψ_ = [10 ∼ 15] *ms*.

We first considered a spine where the influx is confined to the upper pole region of the head. (Figure 6a-e). In this representative case (such as *ρ* = 0.9, *e*_*o*_ = 0.8, and *e*_*i*_ = 0.1), there is a significant localization of high concentration of C close to the pole lasting for more than 20 *ms* (Figure 6b-d). The gradient extends from the top to the bottom of the spine head with the spine apparatus acting as a geometric barrier for diffusion of C. The lifetime of the gradient depends on the geometrical parameters in a similar way as the uniform influx BCs, as shown by the surfaces *τ*_Ψ_(*ρ*, *e*_*i*_, *e*_*o*_) in Figure 6e.

Furthermore, when the PM influx is localized to the side of the spine (Figure 6f-j), a big ER in a spheroidal*−*like spine (*ρ →* 1 and *e_o_ →* 1) still represents a geometrical barrier against the diffusion of *C*. On the other hand, the lifetime of the gradient (Figure 6j) appears to be much shorter (*τ*_Ψ_ = [8 ∼ 15] ms) because the influx is localized closer to the neck, providing a shorter path to the efflux boundary. From these simulations, we conclude that a localized influx on the side of the spine reduces the dependence of *τ*_Ψ_ to the geometrical parameters *ρ* and *e*_*o*_ when compared with the case of influx localized on the pole of the spine or that of uniformly distributed influx (see Figure 6e, j, and Figure 5f).

### 3.5 Mushroom morphology, localization of influx and variation of the size of the spine

Until now we have considered spheroidal geometry with a fixed spine size. Depending on the brain region, spine type, and species the dendritic spine dimensions vary (0.3 ∼ 1.5 *μm* in diameter) [12, 13]. Furthermore, Bourne and Harris observed that heads with diameters bigger than 0.6 *μm* have a mushroom morphology [37]. Therefore, we investigated how the size and shape of the spine head and of the ER (*i.e.* varying *R*_*o*_, *e*_*o*_, *ρ*, and *e*_*i*_), affect the spatio-temporal dynamic of second messengers in idealized mushroom-like geometry with localized influx and efflux in the pole and neck region, respectively (see Figure 7a).

**Figure 7:**
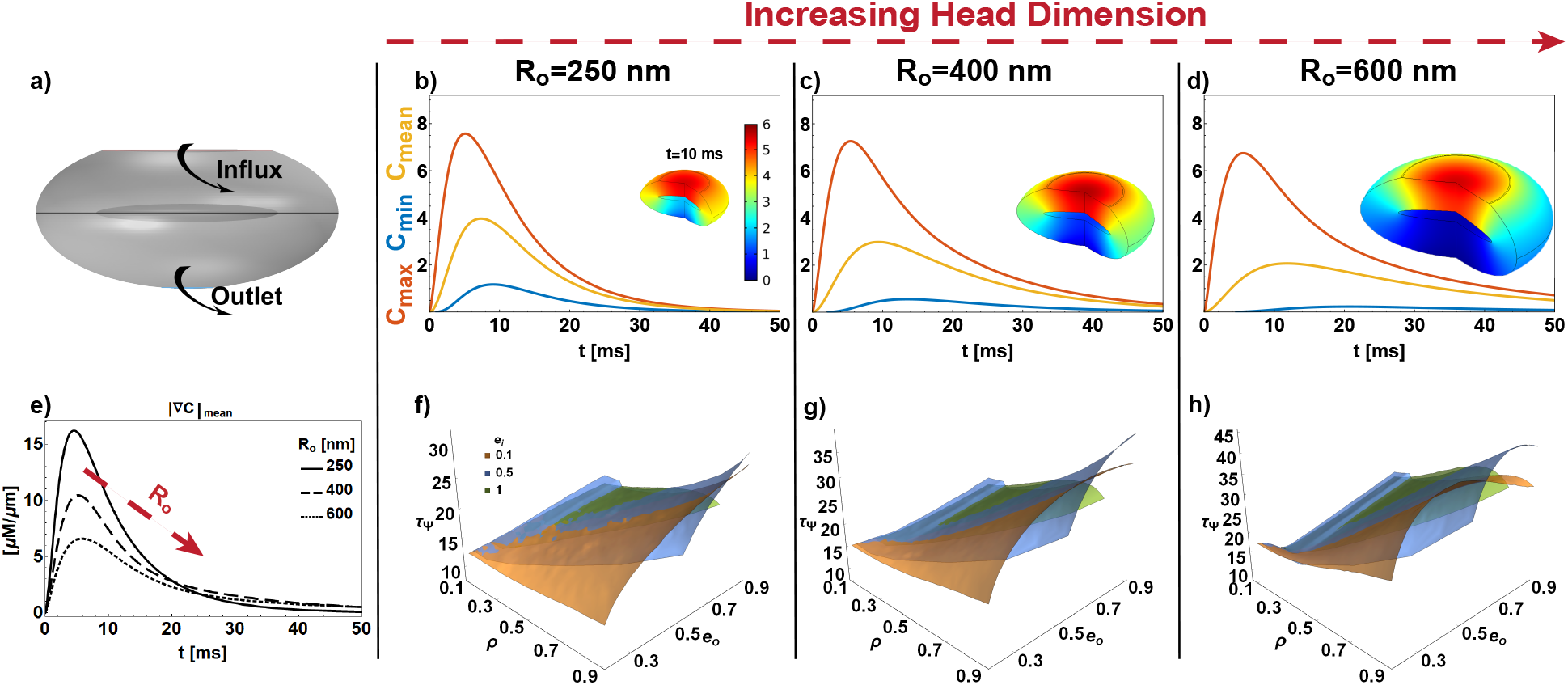
Mushroom-like morphology and spine size effect. In the schematic a) we analyzed a localized pole influx in an idealized mushroom-like morphology with an outlet, considering the geometrical parameters *ρ* = 0.6, *e*_*o*_ = 0.5, and *e*_*i*_ = 0.1. All the simulations shown use *D* = 10 *μm*^2^/*s* and *K*_*N*_ = 1 *μm/s*. The time evolution of *C*_*max*_, *C*_*min*_, and *C*_*mean*_ (b for *R*_*o*_ = 250 nm, c *R*_*o*_ = 400 nm, and d *R*_*o*_ = 600 nm) and the distribution of concentration at a given time (see insets) are not strongly affected by the increasing of *R*_*o*_. Instead, the time evolutions of the gradient (in e)) have lower peak values and longer lifetime (f-g)) as the spine dimension increases (*τ*_Ψ_ up to 45 *ms* in h).

We first maintained a fixed shape (Figure 7a: *ρ* = 0.9, *e*_*o*_ = 0.8 and *e*_*i*_ = 0.1) and varied the spine dimension (*R*_*o*_ = 250, 400, and 600 *nm*). The increase in size slightly affects the spatial distribution and the time course of *C*_*max*_, *C*_*min*_, and *C*_*mean*_ (see Figure 7 and insets). This because the PSD areas scale with the spine size [12]. Furthermore, the higher available volume reduced the peak value of the gradient but simultaneously elongated its lifetime (Figure 7e). In fact, the surfaces of *τ*_Ψ_ as a function of *ρ*, *e*_*i*_, and *e*_*o*_ for the three spine dimensions showed that the lifetime of the gradient increased up to 45 *ms* (Figure 7f-h). Furthermore, from the simulations we noticed that the increase in dimension induced a more complex nonlinear dependence of *τ*_Ψ_ to *ρ* and *e*_*o*_. In fact, we observed a notable non-monotonic trend where *e*_*o*_ increases, a trend that cannot be traced with classical diffusion lifetime alone (compare Figure 5g and Figure 7h).

Finally, comparing between the mushroom-like and spheroidal geometries, both with a localized influx in the pole of the spine (Fig. 6 and Figure 7) we noticed that spatial distribution and the *τ*_Ψ_ surfaces are in very good agreement. Thus, all the geometrical principles discussed for the oblate spheroidal geometries hold for the idealized mushroom-like case.

### 3.6 Nonlinear reaction kinetics affects the spatiotemporal dynamics of C in the long timescale

Thus far, we have only considered linear kinetics of second messengers decay within the domain that could be related to the binding of second messengers to buffers. However, signal transduction events and transport via membranes are often nonlinear, particularly in the dendritic spine [33, 83–85]. We extended our model to investigate whether nonlinear kinetics would change the effect of geometry and BCs. To do this, we modified the reaction-diffusion equation in Equation (1) to include Michaelis–Menten [86] kinetics as follows

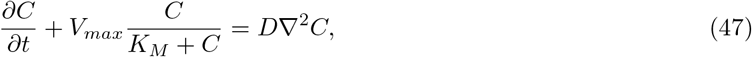

where the maximum rate is *V*_*max*_ = *C_max_/τ* and *K*_*M*_ is the Michaelis-Menten constant.

We performed the simulations with *C*_*max*_ = 4 *μ*M, *τ* = 50 ms, and *K*_*M*_ = 2 *μ*M, chosen for sake of representation and compared the results obtained with linear reaction kinetics (Equation (1)). The comparison of the case previously presented (Figure 8a-c) showed that the complexity introduced with a nonlinear reaction does not affect the spatiotemporal dynamics of second messengers in the short timescale (*t <* 10 ms), but affects the long-term dynamics showing a decay with different slopes. Furthermore, the localization and value of the maximum concentration do not depend on the kinetics, but rather are governed by the geometry and BCs. The kinetics affects the dynamics when a well-mixed distribution is already reached in all the cases, exhibiting different decay rates.

**Figure 8:**
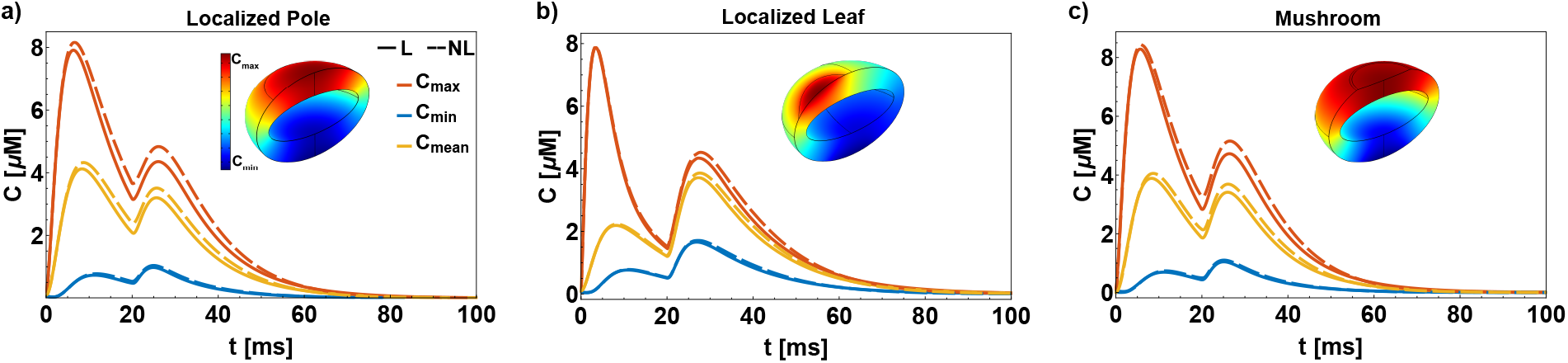
Comparison between linear (solid lines) and nonlinear (dashed lines) kinetics for the ellipsoidal geometry with localized pole a), localized leaf b), and mushroom-like morphology c). All the simulations shown use *D* = 10 *μm*^2^/*s*, *R*_*o*_ = 250 *nm*, and *K*_*N*_ = 1 *μm/s*. The nonlinear nature of the kinetics does not change the localization of the maximum (in red), minimum (in blue), mean (in yellow), and the lifetime of the localization. However, the nonlinear kinetics affect the long timescale dynamics (*t >* 10 ms) resulting in decay with a different slope.

## 4 Discussion

Recent experimental observations have presented detailed, high-resolution images of the architecture of dendritic spines highlighting a complex internal organization [40, 79–82]. Such observations serve to highlight the role of geometry and spatial features in cellular phenomena. In this work, we used a general framework to study the effect of spine geometry including the internal organization in an idealized mathematical model with the goal of identifying some governing principles that regulate the spatio-temporal dynamics of second messengers. To do this, we developed and analyzed a general mathematical model, in which, a reaction-diffusion partial differential equation (PDE) with time-dependent mixed boundary conditions (BCs) were analytically and numerically solved.

We arrive at the following conclusions form this work. First, the lifetime of second messengers gradients in dendritic spines depends on the intrinsic coupling between geometry of the spines and boundary conditions. These boundary conditions reflect the signaling events that take place at the membrane. Numerical simulations demonstrated how the shape of the spine governs the transient localization of peak concentration of second messengers. The lifetime of this localization depends nonlinearly on the geometry. Furthermore, we also showed that localized BCs, confined to a portion of the boundary, reduce the effect of the presence of a big and spheroidal-like ER if the influx is on the side of the spine head. Analytical investigation (Section 3.1) showed that geometry dictates the specific kind of harmonic function for the spatial distribution of second messengers. The temporal dynamics in the long time is governed by the kinetics of signaling reaction in the cytoplasm. However, this separation of temporal and spatial effects is not straightforward. The time-dependent BCs (Equation (2)) represent both the kinetics of the membrane reactions and the curvature of the boundary (in the 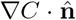 term). Therefore, time-dependent BCs represent the coupling between shape and kinetics (Figure 9).

**Figure 9:**
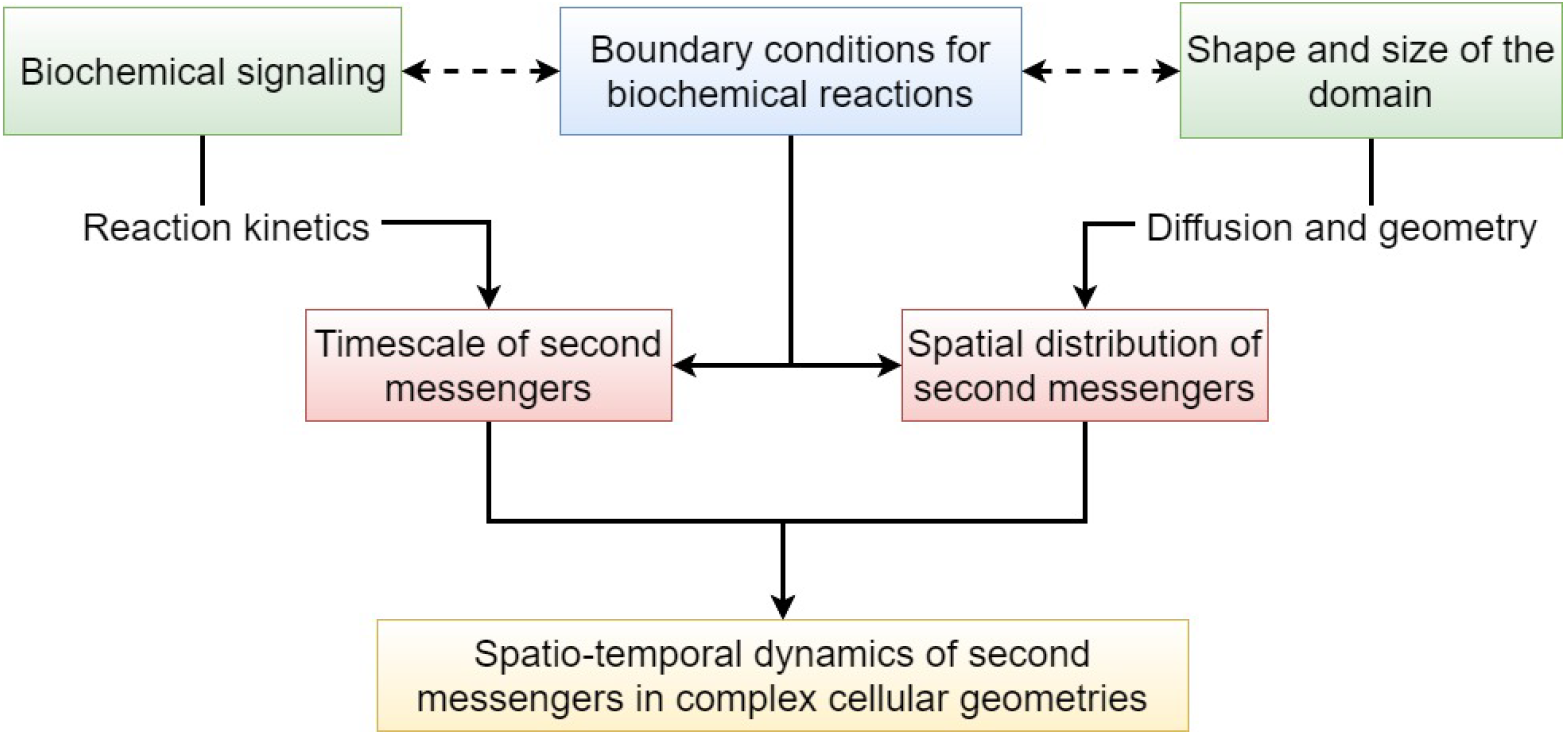
Geometric principles of second messengers dynamics: The spatio-temporal dynamics of second messengers are affected by the interplay of shape and size, biochemical signaling, and membrane reactions through boundary conditions. Our study showed that the amplitudes of the harmonic functions found in the analytical solution and dictated by the geometry, intricately depend on the boundary condition in the short-timescale and on the reaction kinetics on long timescale.

Second, localization of the fluxes plays an important role in governing the spatiotemporal dynamics of second messengers. The lifetime of the gradient is affected by the pole vs. side localization of the spine head suggesting that the localization of the PSD plays a crucial role in spine signaling. Third, the organelle membrane plays two roles. One is to act as a diffusion barrier and the second is to act as source or sink of BCs. An emerging idea in shape regulation of signaling is the role played by organelles such as the endoplasmic reticulum and nucleus [53, 87]. In the case of spines, the spine apparatus is thought to play a critical role in governing synaptic plasticity [42–48]. We find that the relative organization of the two membranes, PM and the ER membrane, affect the geometric landscape through both shape and boundary condition effects. We also find that the organelles can act as a physical barrier to diffusion extending the lifetime of the gradient. And finally, at short timescales, the nature of the kinetics in the cytoplasm does not alter our conclusions but kinetics play an important role in the long timescale especially in coupled cascades [3, 5, 88, 89]. These predictions are applicable not just to spines but also to cells in general.

Even though our model is simplified, it allows us to identify some common principles by which geometry can be used to alter timescales of signal transduction. The notion that shape alone can affect signal transduction (Figures 4 and 5) is a principle that is now being well-accepted in the literature [3–5, 7–10] and we now extend this idea to signaling subcompartments such as dendritic spines.

Despite the relatively simple model presented here, we found that the spatial dependence of second messengers at short timescales was nonintuitive and exhibited a complex dependence on geometry. One potential design principle that we have identified here is that dendritic spines may be able to ensure rapid and robust signal propagation from the head toward the neck and thus toward the dentritic shaft by combining flattened head shape with very small or absence of ER, especially if the influx is localized at the pole of the spine. On the contrary, if the goal is to localize second messengers for a longer time, such as to promote synaptic plasticity, a design that allows the growth of bigger ER in a spheroidal-shaped head would help to ensure longer lifetime of the gradient. Future efforts need to consider the dynamics of both the spine and the spine apparatus during structural plasticity to incorporate mechanochemical effects.

Based on these insights, the next steps in spine systems biology can focus on specific signal transduction pathways and use reconstructions of realistic geometries to identify how the simple mathematics presented here translate into computational biology. Additionally, experimentally advances in localized imaging of second messengers molecules will be necessary to test and validate the predictions made by computational modeling. These combined efforts will enable us to extend these simple models to biologically relevant processes.

## Acknowledgments

We thank Dr. Lucas Stolerman, Dr. Donya Ohadi K. M., Ms. Miriam Bell, and Mr. Arijit Mahapatra for the feedback on the manuscript. AC, TMB, TJS, and PR would also like to thank the AFOSR (grant number FA9550-18-1-0051) and RI the NIH (grant number GM072853) for funding support.

## Author Contributions

P.R. designed the study. A.C. developed the analyses and performed the simulations. All authors discussed the results, wrote the text and contributed to the final version of the manuscript.

